# The Broad Role of Nkx3.2 in the Development of the Zebrafish Axial Skeleton

**DOI:** 10.1101/2020.12.30.424496

**Authors:** Laura Waldmann, Jake Leyhr, Hanqing Zhang, Caroline Öhman-Mägi, Amin Allalou, Tatjana Haitina

## Abstract

The transcription factor Nkx3.2 (Bapx1) is an important chondrocyte maturation inhibitor. Previous *Nkx3.2* knock-down and overexpression studies in non-mammalian gnathostomes have focused on its role in primary jaw joint development, while little is known about the function of this gene in broader skeletal development. We generated CRISPR/Cas9 knockout of *nkx3.2* in zebrafish and applied a range of techniques to characterize skeletal phenotypes at developmental stages from larva to adult, revealing fusions in bones of the occiput, the loss or deformation of bony elements derived from basiventral cartilages of the vertebrae, and an increased length of the proximal radials of the dorsal and anal fins. These phenotypes are reminiscent of *Nkx3.2* knockout phenotypes in mammals, suggesting that the function of this gene in axial skeletal development is ancestral to osteichthyans. Our results highlight the broad role of *nkx3.2* in zebrafish skeletal development and its context-specific functions in different skeletal elements.

## Introduction

NK3 homeobox 2 (Nkx3.2, Bapx1) is an evolutionarily conserved gene encoding a homeodomain-containing transcription factor that is involved in cartilage growth and differentiation in gnathostomes. It was first described in Drosophila (*bagpipe, bap*), where it plays a major role in the visceral mesoderm during the formation of the midgut musculature (Azpiazu and Frasch, 1993). During vertebrate evolution *Nkx3.2* expression was incorporated into the intermediate domain of the first pharyngeal arch. This event has been proposed to be crucial for jaw joint formation during the transition from jawless to jawed vertebrates (Cerny et al., 2010). Jawed vertebrates like the zebrafish, frog, and chicken display a focal expression of *Nkx3.2* between Meckel’s and palatoquadrate cartilages of the first pharyngeal arch skeleton in contrast to the jawless lamprey, which shows more diffuse expression (Kuraku et al., 2010; Miller et al., 2003; Square et al., 2015; Wilson and Tucker, 2004). During development, ventrally migrating cranial neural crest cells form the first pharyngeal arch skeleton. This process is controlled by signalling molecules like endothelin-1 (Edn1), which positively regulates Nkx3.2 (Miller et al., 2003; Nair et al., 2007). Previous studies showed that Nkx3.2 is an essential factor for primary jaw joint development, as the loss of expression in zebrafish, frog, and chick leads to a failure in jaw joint formation accompanied by the fusion of the joint articulating cartilage elements: Meckel’s cartilage and the palatoquadrate (Lukas and Olsson, 2018a; Miller et al., 2003; Wilson and Tucker, 2004). Overexpression of *nkx3.2* (*bapx1*) in amphibians induces the formation of ectopic cartilage elements by introducing additional subdivisions into existing cartilage, clearly showing the joint-promoting effect of this transcription factor (Lukas and Olsson, 2018b).

As a consequence of the primary jaw joint to middle ear transition during the course of mammalian evolution (Luo, 2011; Anthwal *et al*., 2013), *Nkx3.2* is expressed within the middle ear-associated bones of the tympanic ring and gonium as well as in the incudomalleolar joint in mammals (Tucker et al., 2004). Further expression analysis in mouse embryos showed *Nkx3.2* expression in the developing vertebrae and the cartilaginous condensations of the developing limbs (Tribioli et al., 1997). Mouse embryos deficient in *Nkx3.2* display hypoplasia in the tympanic ring, the absence of the gonium, a size reduction of cranial occipital bones such as the basioccipital and basisphenoid, and finally the loss of the supraoccipital bone and vertebral ossification centres (Lettice et al., 1999; Tribioli and Lufkin, 1999; Tucker et al., 2004). Various studies describe Nkx3.2 as a chondrocyte maturation inhibitor during chondrogenesis (Caron et al., 2013; Lengner et al., 2005; Provot et al., 2006; Yamashita et al., 2009). In chicken and mouse long bone development, Nkx3.2 can repress the chondrocyte maturation factor Runx2 during endochondral ossification and thus maintaining the chondrocytes in an immature state (Provot et al., 2006). In humans the homozygote mutation in NKX3.2 gene leads to a spondylo-megaepiphyseal-metaphyseal dysplasia (SMMD), a rare skeletal disease (Hellemans et al., 2009; Simsek-Kiper et al., 2019). The patients suffer from, among other symptoms, a short stature, stiff neck and trunk, and defects in vertebral ossification (Agarwal et al., 2003; Hellemans et al., 2009; Silverman and Reiley, 1985; Simon et al., 2012; Simsek-Kiper et al., 2019), similar to what was observed in mouse knockout mutants. These data clearly indicate a role of Nkx3.2 in the mammalian axial skeleton beyond just the middle ear that is homologous to the non-mammalian primary jaw joint, and yet the function of this gene in the axial skeleton of non-mammals has not been investigated in detail. In zebrafish embryos and juveniles *nkx3.2* expression can be detected in the anterior notochord, vertebrae, and median proximal radials (Arnold et al., 2015; Crotwell and Mabee, 2007).

In this study, we present a comprehensive investigation of the function of *nkx3.2* in zebrafish skeletal development by generating a CRISPR/Cas9 induced mutant line and characterizing larval, juvenile, and adult phenotypes. Our results confirm and elaborate on the primary jaw joint loss previously reported in knockdown studies and we describe novel axial phenotypes in the occiput, Weberian apparatus, rib-bearing vertebrae, and median fins, pushing back the likely origin of axial functions of Nkx3.2 to the osteichthyan stem group.

## Results

### Zebrafish nkx3.2^uu2803/uu2803^ line generated with CRISPR/Cas9 survives to adulthood and has open mouth phenotype

Clustered Regulatory Interspaced Short Palindromic (CRISPR)/CRISPR-associated protein 9 (Cas9) was used to generate *nkx3.2* mutant line to analyse both embryonic and adult mutant phenotypes. We generated a mutant line with a 7 bp deletion in *nkx3.2* exon 1 (c.286_292 del, p.Lys95*), causing a frameshift, which resulted in a premature stop codon that shortens the peptide sequence to 95 amino acids, compared to 245 amino acids in the wild-type (Figure 1A-B). The DNA-binding homeodomain of Nkx3.2 is absent in this shortened protein.

**Figure 1 –.**
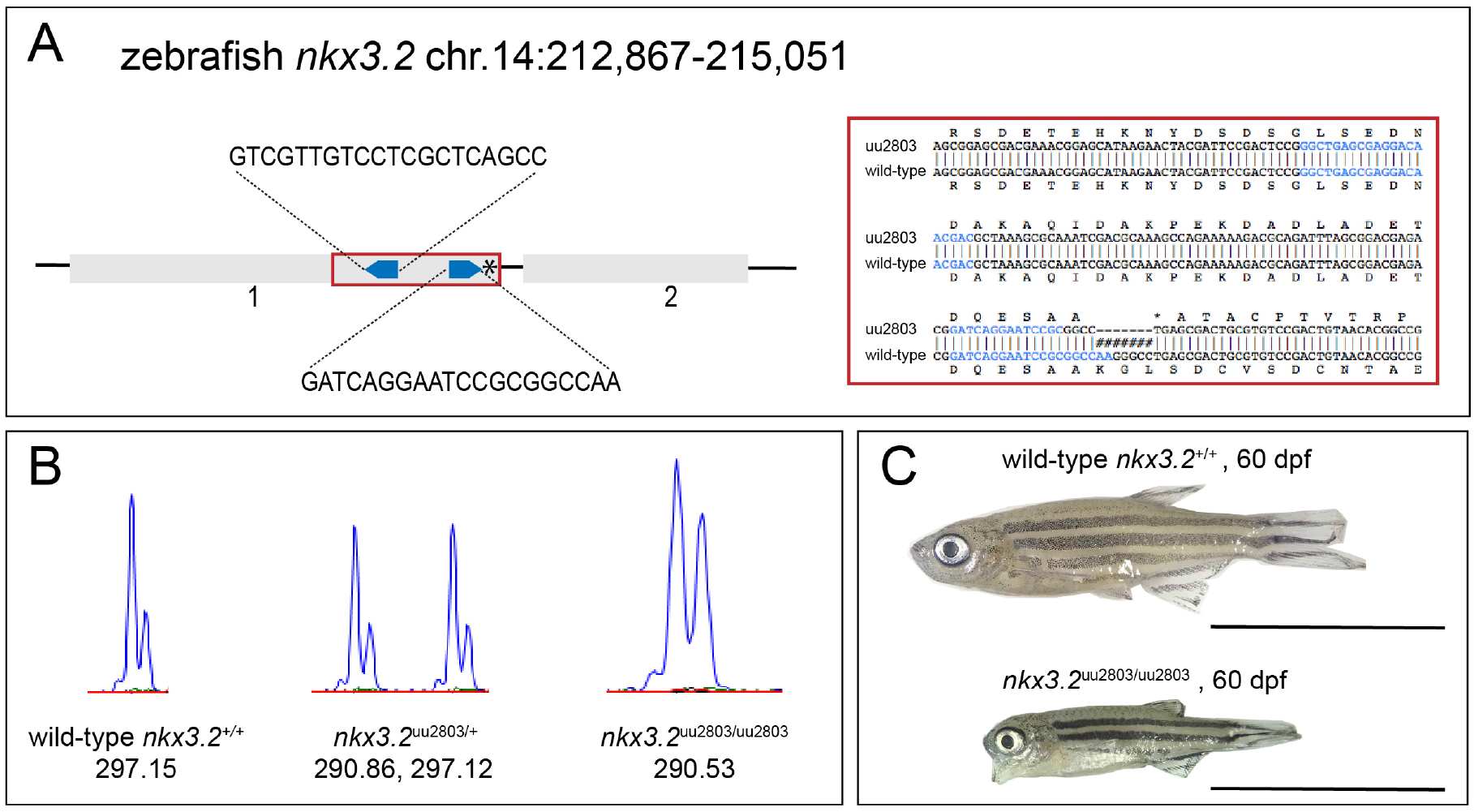
Zebrafish *nkx3.2* knockout generated with CRISPR/Cas9. (**A**) Schematic diagram of zebrafish *nkx3.2* exons one and two marked by grey rectangles with two CRISPR target sides marked by blue arrowheads. The 7 bp deletion in the first exon is marked by the asterisk in exon 1. The deletion causes an early a stop codon after 95 amino acids (confirmed by sequencing) (**B**) Wild-type, heterozygous *nkx3.2*^uu2803/+^ and homozygous *nkx3.2*^uu2803/uu2803^ fish were analysed by fragment length analysis. The wild-type displayed one peak (297,15), *nkx3.2*^uu2803/+^ fish one wild-type (297,12) and one mutant peak (290,86) and *nkx3.2*^uu2803/uu2803^ fish one mutant peak (290,53). (**C**) At 60 dpf *nkx3.2*^uu2803/uu2803^ fish display a prominent fixed open mouth phenotype. Scale bars: 1 cm.

Heterozygous embryos (*nkx3.2*^uu2803/+^) displayed no morphological differences compared to wild-type (*nkx3.2*^+/+^) embryos as long as observed (data not shown). Homozygous mutant embryos displaying morphological differences were generated by crossing two heterozygous *nkx3.2*^uu2803/+^ adult zebrafish. Homozygous mutants (*nkx3.2*^uu2803/uu2803^) were able to survive to adulthood and displayed a prominent fixed open mouth phenotype (Figure 1C). Adult homozygous mutants displayed differences in size relative to adult wild-type, although it was difficult to determine whether this was due to the mutation or a consequence of the fixed open mouth phenotype effect on feeding (Figure S1).

### Confocal live imaging displays fusion of Meckel’s cartilage with palatoquadrate in nkx3.2^uu2803/uu2803^ zebrafish embryo and larvae

In order to follow the effects of *nkx3.2* CRISPR/Cas9 induced knockout within the first mandibular arch during development, confocal live imaging was performed. To visualise the pharyngeal arches, *Tg*(*sox10:egfp*) line labelling neural crest-derived cells was used. Clear phenotypic differences in the jaw joint-forming region were detectable from 3 dpf onwards (Figure 2). The jaw joint was lost in *nkx3.2*^uu2803/uu2803^ fish as Meckel’s cartilage and the palatoquadrate were fused (Figure 2 E-H’) and the retroarticular process (RAP) was missing at the posteroventral tip of Meckel’s cartilage (Figure 2). Knockout of *nkx3.2* furthermore resulted in an unorganised cell-mass of small rounded chondrocytes in the area where the jaw joint is normally formed at 3 and 5 dpf, whereas chondrocytes of Meckel’s cartilage and the palatoquadrate more distal to the jaw joint-forming region were elongated and displayed typical stacking (Figure 2 E, E’, F, F’). By 7 dpf, chondrocytes at the fusion site acquired a more elongated shape and began to align with the stacked chondrocytes of Meckel’s cartilage and the palatoquadrate (Figure 2 G, G’). At 14 dpf, all chondrocytes at the fusion site were elongated and aligned completely with the adjacent chondrocytes (Figure 2 H, H’).

**Figure 2 –.**
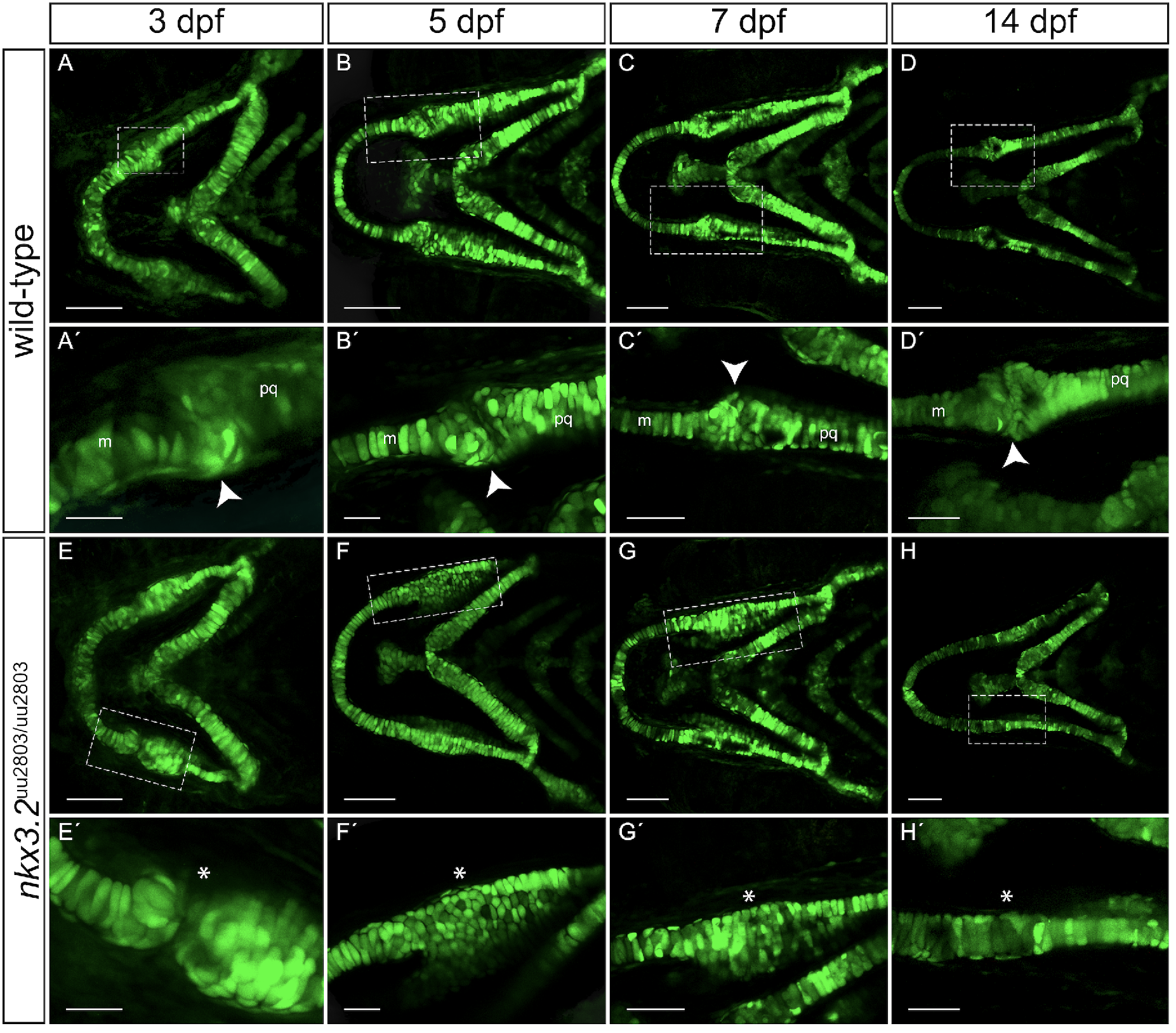
Larval development of wild-type and *nkx3.2^uu2803/uu2803^* jaw joints. (**A-H’**) Maximum projections of confocal live imaging Z stacks acquired from ventral side of wildtype zebrafish head at 3 dpf (**A, A’**), 5 dpf (**B, B’**), 7dpf (**C, C’**), 14 dpf (**D, D’**) and *nkx3.2^uu2803/uu2803^* zebrafish head at 3 dpf (**E, E’**), 5 dpf (**F, F’**), 7 dpf (**G, G’**), 14 dpf (**H, H’**) in *Tg*(*soxl0:egfp*) background. (**A-D**) The jaw joint in wild type zebrafish (dashed box) is magnified in **A’-D’**. The retroarticular process is visible from 3 dpf onwards, marked by white arrowhead. Chondrocytes of Meckel’s cartilage and palatoquadrate align in stacks. Posterior Meckel’s cartilage and anterior palatoquadrate articulate the jaw joint. (**E-H**) Fusion of jaw joint articulating elements in *nkx3.2* mutant fish. The fusion is magnified in (**E’-H’)** and indicated by asterisks. (**E’-F’**) 3 dpf and 5 dpf *nkx3.2* mutant embryos display unorganised and rounded cells in the area where the joint would normally form. (**G’**) 7 dpf *nkx3.2* mutant larvae display elongated cells in the fused area, which starts to align in stacks. (**H’**) 14 dpf *nkx3.2* mutant zebrafish display elongated cells that align with adjacent Meckel’s cartilage and palatoquadrate. m – Meckel’s cartilage, pq – palatoquadrate. Scale bars: 100 μm (**A-H**), 25 μm in (**A’-H’**).

### Histological analysis of nkx3.2^uu2803/uu2803^ zebrafish shows chondrocyte alignment and hypertrophy at the jaw joint fusion site

Histological sections were prepared to further analyse the chondrocyte arrangement within the first mandibular arch in *nkx3.2* mutant larval, juvenile, and adult zebrafish. A joint gap between Meckel’s cartilage and the palatoquadrate is visible in wild-type zebrafish at 14 dpf.

Articular chondrocytes lining the joint cavity and hypertrophic chondrocytes forming the articulating elements were present (Fig. 3 A and A’). In *nkx3.2* mutant larvae at 14 dpf, Meckel’s cartilage and the palatoquadrate were not separated but fused – the jaw joint was absent. Chondrocytes within the fused element were hypertrophic and aligned. At the presumptive fusion site, the element appeared to be increased in width caused by piled-up rows of aligned chondrocytes (Fig. 3 B and B’). The exact fusion point was difficult to determine in both 30 dpf and 90 dpf mutant fish but apart from the fusion and its phenotypic consequences, ossification seemed not to be affected. Articular chondrocytes lining the articulating tips of the articular and quadrate were consequently absent in *nkx3.2* mutant juvenile fish at 30 dpf. Chondrocytes within the fused element underwent hypertrophy as in wild-type fish (Fig. 3 C-D’). By 90 dpf, ossification of the fused articular and quadrate was completed, visible by the presence of adipose tissue inside the bones in both wild-type and mutant adult fish (Fig. 3 E-F’).

**Figure 3 –.**
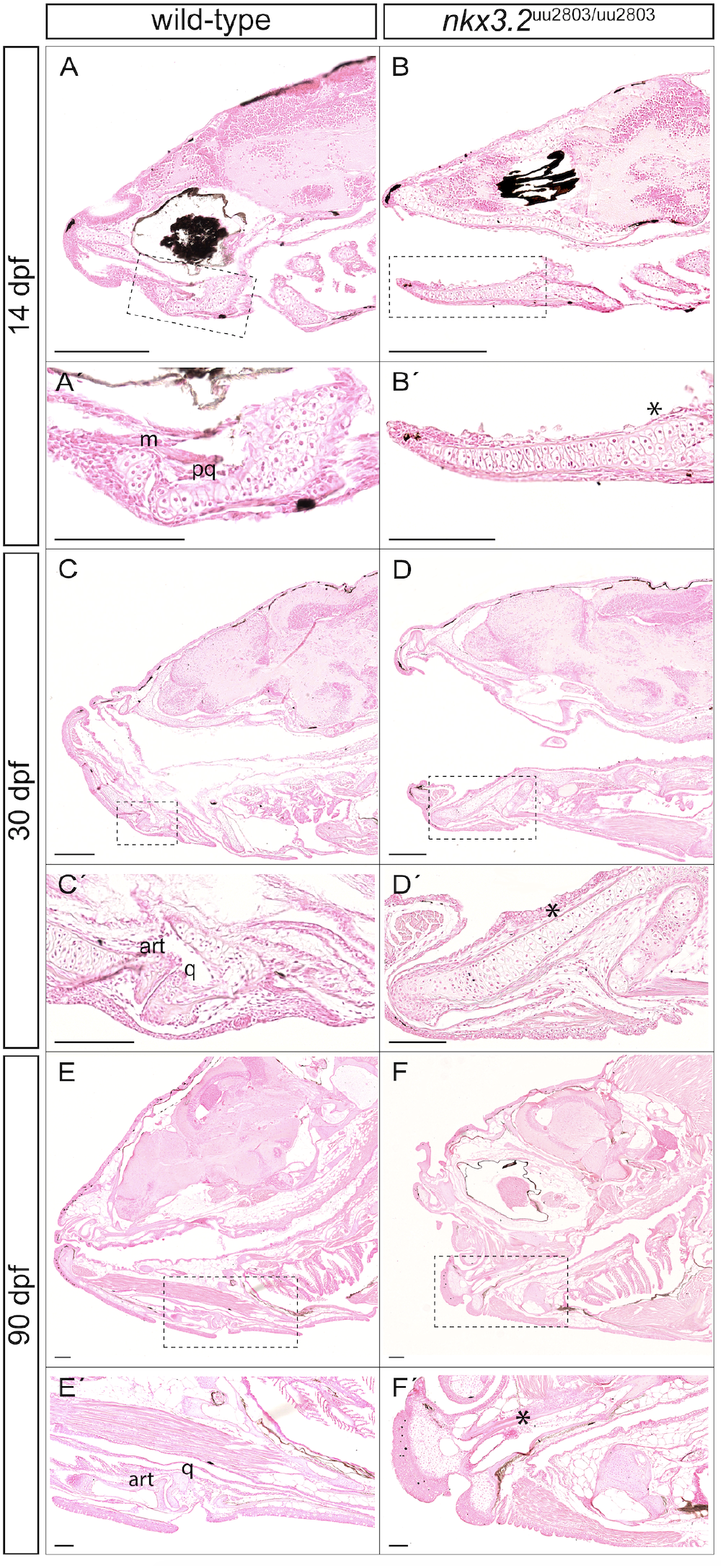
Histological analysis reveals loss of jaw joint without affected chondrocyte hypertrophy in the first pharyngeal arch of *nkx3.2^uu2803/uu2803^* zebrafish. Sagittal sections of wild-type 14 dpf (**A, A’**), 30 dpf (**C, C’**) and 90 dpf (**E, E’**) and *nkx3.2* mutant 14 dpf (**B, B’**) 30 dpf (**D, D’**) and 90 dpf (**F, F’**) zebrafish stained with Nuclear Red Stain. (**A, A’, C, C’, E, E’**) Stained histology sections displaying a normal jaw joint development between Meckel’s cartilage (m) and palatoquadrate (pq), respectively articular (art) and quadrate (q) in wildtype fish. (**B, B’, D, D’, F, F’**) *nkx3.2* mutant fish do not display a jaw joint. Chondrocyte maturation and ossification seem not to be affected in *nkx3.2^uu2803/uu2803^* besides the absence of joint-typical cells lining the articulating elements. Dashed box in (**A-F**) marks the magnified region in (**A’-F’)**. Asterisk mark the jaw joint fusion. m – Meckel’s cartilage, pq – palatoquadrate, art – articular, q – quadrate. Scale bars: 200μm (**A-F**), 100μm (**A’-F’**).

### Optical projection tomography reveals morphological changes in the head of nkx3.2^uu2803/uu2803^ larvae

In order to characterise the larval *nkx3.2* mutant phenotype in greater detail, we used optical projection tomography (OPT) on 5 dpf cartilage-stained larvae to reconstruct 3D models of wild-type and mutant cartilage morphology. Multiple wild-type (n=10) and mutant (n=11) 3D models were overlaid and combined to produce an average wild-type morphology (Figure 4A’ D, G) and an average *nkx3.2* mutant morphology (Figure 4B, E, H). Overlaying these grouped 3D reconstructions allowed the calculation of voxels that were statistically more or less intense in the mutant group, corresponding to locations with more or less cartilage present, respectively. This analysis was robust to false positives (Figure S2).

**Figure 4 –.**
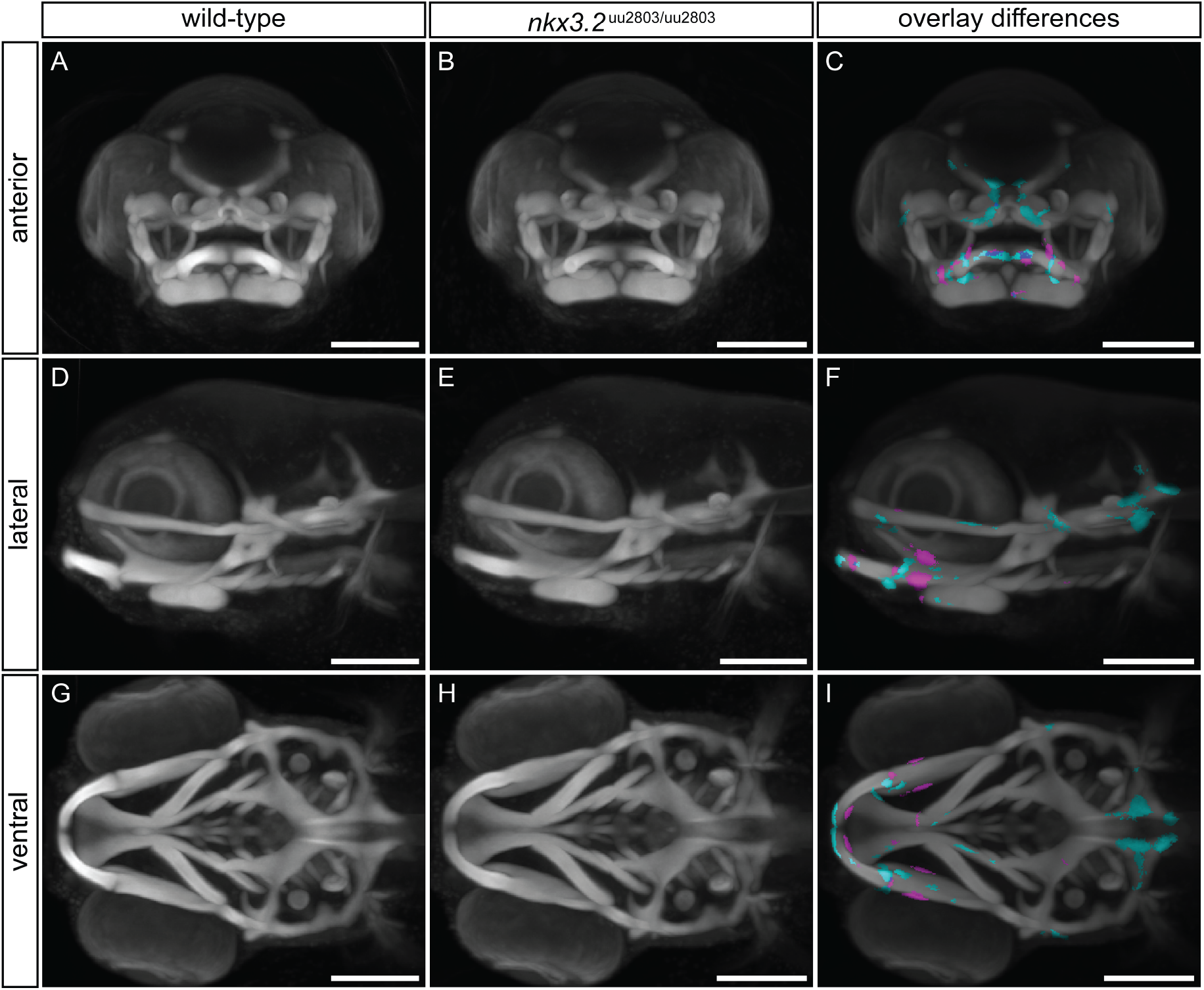
Optical projection tomography of cartilage-stained wild-type and *nkx3.2^uu2803/uu2803^* larvae reveals subtle phenotypic changes. (**A, D, G**) Maximum projection of 5 dpf wild-type group (n=10). (**B, E, H**) Maximum projection of 5 dpf *nkx3.2* mutant (n=11) group. (**C, F, I**) Maximum projection of both groups with coloured voxels representing voxels with statistically significant (p<2.5×10^-4^) differences in intensity. Cyan shows voxels with lower intensity and magenta shows voxels with higher intensity in *nkx3.2* mutant group. Scale bars: 300μm.

The *nkx3.2* mutant group showed significantly higher intensity of the cartilage labelling at the jaw joint region (magenta in Figure 4F, I) consistent with the presence of fused cartilage in mutants compared to the jaw joint gap in wild-type larvae. Cyan voxels ventral to the jaw joint in Figure 4F indicate the absence of the RAP. The palatoquadrate displayed significantly higher intensity in *nkx3.2* mutant group consistent with the increased thickness of this element (magenta in Figure 4C, F, I). The anterior part of Meckel’s cartilage displayed an increased posterior intensity and decreased anterior intensity in the *nkx3.2* mutant group indicating a subtle change in the shape of Meckel’s cartilage (Figure 4I). Interestingly, this analysis also revealed significantly reduced cartilage staining signal in the posterior part of the head, around the otic capsule (Figure 4 F, I). The mutant group clearly showed the shorter parts of the auditory capsule and thinner parachordal changing the shape of the notochord insertion, compared to the wild-type group (Figure 4F, I).

### Cartilage and bone staining analysis of wild-type and nkx3.2^uu2803/uu2803^ larval, juvenile and adult zebrafish

Skeletal staining of larval, juvenile and adult *nkx3.2* mutant zebrafish was performed to analyse both cartilage and bone abnormalities in comparison to wild type at a greater range of developmental stages. The loss of the jaw joint caused by the fusion of Meckel’s cartilage and palatoquadrate was clearly visible from 5 dpf onwards and was most recognizable by the absence of the RAP (Figure 5). The resulting open mouth phenotype could be observed from 9 dpf onwards (Figure 5D, F, H). No other cartilage phenotypes within the pharyngeal arches could be detected at any developmental stage, consistent with the OPT results at 5 dpf in Figure 4. Ossification of the dentary, articular, and quadrate appeared to take place normally in *nkx3.2* mutants at 30 dpf (Figure 5H’) when compared to wild-type fish (Figure 5G’), although the absence of the cartilaginous jaw joint resulted in an ossified fusion between the articular and quadrate.

**Figure 5 –.**
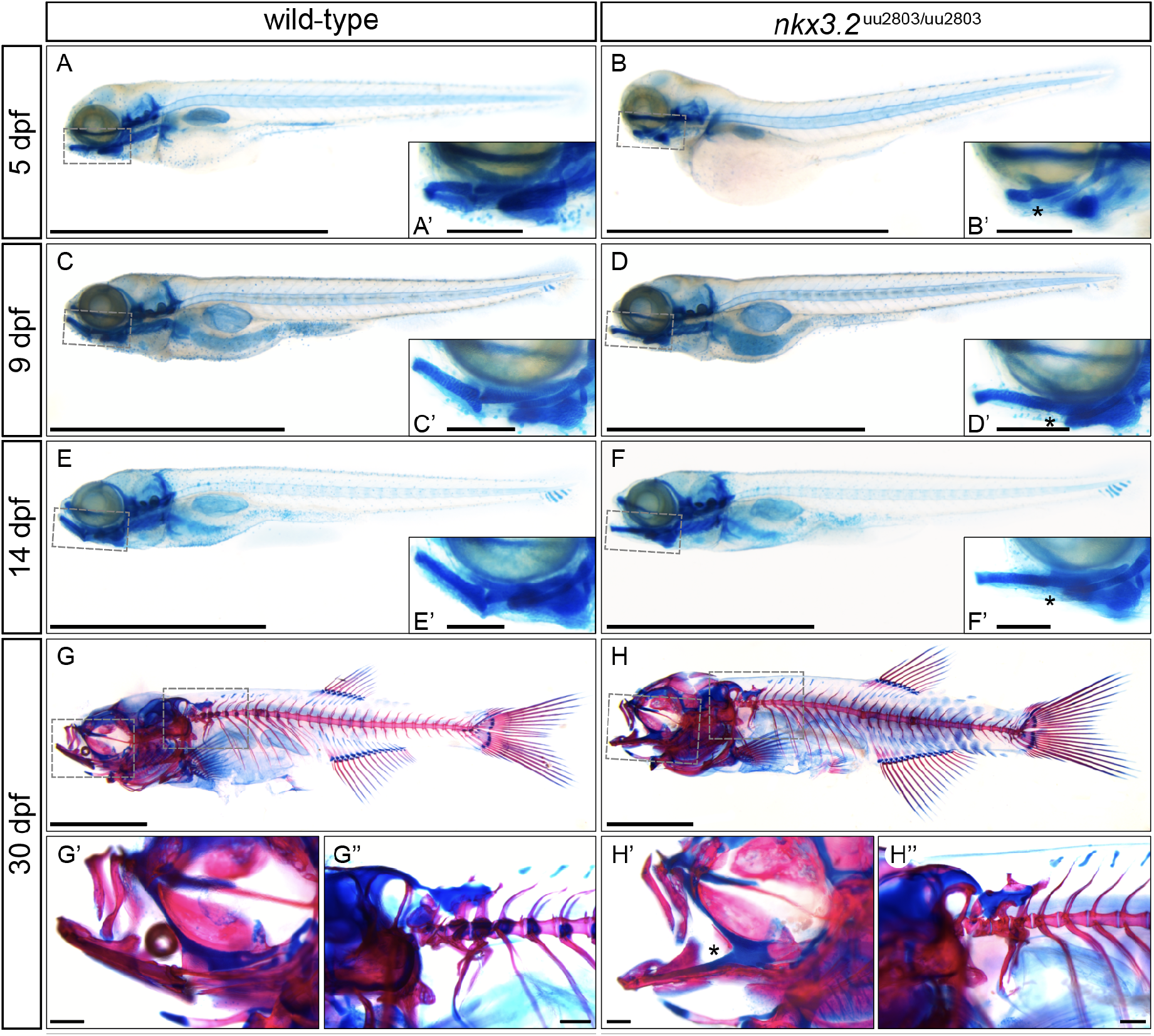
Skeletal staining of wild-type and *nkx3.2^uu2803/uu2803^* zebrafish. (**A-F**) lateral views of cartilage-stained wild-type and *nkx3.2* mutants from 5-14 dpf. (**G-H**) lateral views of cartilage- and bone-stained wild-type and *nkx3.2* mutants at 30 dpf. Boxes in (**A-H**) indicate the zoomed-in regions in the insets or zoomed-in panels (**G’**, **G’’**, **H’**, **H’’**). Asterisks indicate the fusion between Meckel’s cartilage and palatoquadrate in the jaw joint. Scale bars: 2mm (**A-H**), 200μm (**A’-H’’**).

### μCT reveals craniofacial phenotypes in adult nkx3.2^uu2803/uu2803^ zebrafish

In order to examine the axial skeleton of adult zebrafish we performed μCT followed by 3D reconstruction of wild-type and *nkx3.2* mutant zebrafish at 90 dpf (Figure 6). The resting position of the mouth is closed (CM) in wild-type individuals, so to be better able to compare the phenotype of the wild-type to the open-mouthed mutant, μCT scans were also performed on 90 dpf zebrafish with their mouths held open (OM) with a pipette tip. The fusion of the Meckel’s and palatoquadrate cartilages early in the development of *nkx3.2* mutants effectively resulted in a fusion of the articular and quadrate that ossify from these cartilage precursors. This made it extremely challenging to demarcate the articular and quadrate in the *nkx3.2* mutants at 90dpf, which is why the lower jaw was not segmented in red in Figure 6B, E, H, K. As in younger *nkx3.2* mutants (Figure 5D, F, H), the mouth of 90dpf mutants was fixed in an open position, and this resulted in opposing forces being exerted between the lower jaw, the basihyal, and the second pharyngeal arch, the ceratohyal. The outcome of these forces varied between different individuals. Figure 6E shows an individual where the ventral position of the lower jaw appeared to have pushed the basihyal and ceratohyal posteroventrally, resulting in a sharp angle between the anteroventrally-pointing anterior end of the ceratohyal and the posteroventrally-pointing posterior end of the basihyal. Other individuals (not shown) had a relatively normal position of the basihyal and ceratohyal in the mouth compared to the ventrally-positioned lower jaw, resulting in the basihyal partially obstructing the open mouth, its anterior end positioned dorsally to the entire lower jaw. In some cases, the opposing forces exerted by the lower jaw and ceratohyal on either side of the basihyal caused it to bend and ossify into an L-shape.

**Figure 6 –.**
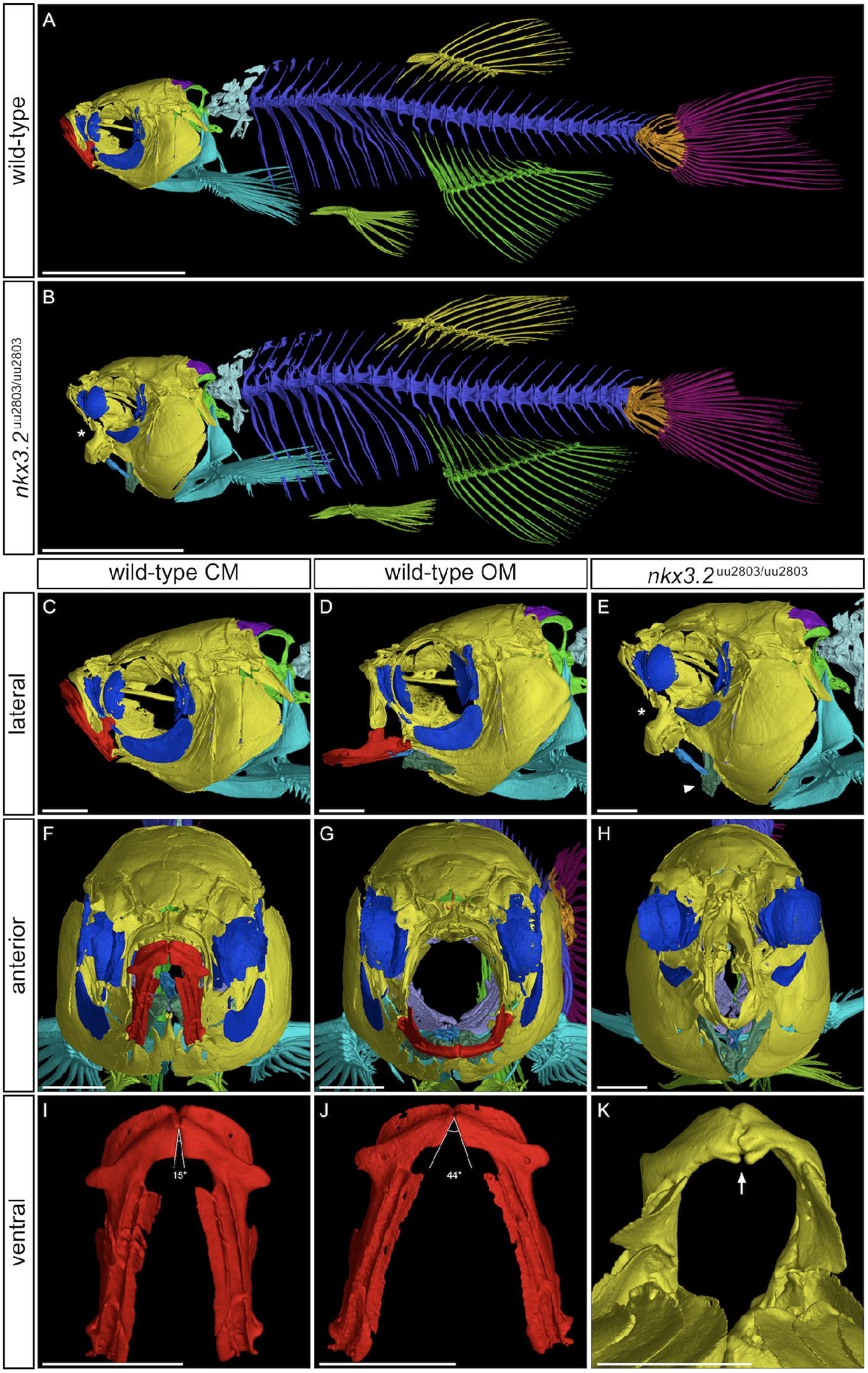
μCT reveals craniofacial phenotypes in adult wild-type and *nkx3.2^uu2803/uu2803^* zebrafish. **(A, B)** lateral view of wild-type and *nkx3.2* mutant zebrafish at 90dpf. **(C, D, E)** lateral view of the head of wild-type with closed mouth (CM), wild-type with open mouth (OM), and *nkx3.2* mutant. **(F, G, H)** Anterior view of wild-type with CM, wild-type with OM, and *nkx3.2* mutant. **(I, J, K)** ventral view of isolated wild-type dentary (CM), wild-type dentary (OM), and *nkx3.2* mutant dentary. Asterisks in **(B, E)** indicate the fused jaw joint phenotype, the arrowhead in **(E)** indicates the downturned ceratohyal phenotype relative to **(D)**. Angles in **(I, J)** show the flexibility of the symphysis in wild-type adults, the arrow in **(K)** indicates the deformed symphysis phenotype in *nkx3.2* mutant. Colour scheme: red – lower jaw; dark blue – infraorbitals, lateral ethmoid, and pterosphenoid; dark green – ceratohyal; blue – basihyal; violet – branchial arches; dark purple – supraoccipital; green – exoccipital and basioccipital; cyan – cleithrum and pectoral fins. Scale bars: 5mm **(A, B)**, 1mm **(C-K)**.

Viewed anteriorly (Figure 6F-H), the face of the mutant appeared “pinched” at the position where the jaw joint would have formed, resulting in a reduced area of the mouth opening and an anteromedial rotation of more posterior structures, most notably the pterosphenoid, lateral ethmoid, and infraorbital 3. There were also impacts on the bones of the upper jaw, the premaxilla and maxilla, which also appeared to be posteriorly compressed into the cranium (note the reduced distance between the lateral ethmoid and maxilla in figure 6E relative to 6C, D) and compressed laterally in line with the “pinched” jaw apparatus. Contrary to Miyashita et al. (2020), we found that the kinethmoid was present, not absent, at both 60 and 90 dpf, and relatively morphologically unchanged relative to wild-type fish (Figure S3).

The cartilaginous symphysis joint between the paired bones of the dentary flexes during feeding in wild-type zebrafish as the jaw is opened and the width of the mouth opening is increased by the lateral flaring of the suspensorium (Gidmark et al., 2012; Westneat, 2006), illustrated in Figure 6I, J. In contrast, the symphysis of *nkx3.2* mutants appeared to be fused (Figure 6K) and was likely inflexible.

### Juvenile and adult nkx3.2^uu2803/uu2803^ zebrafish display loss of basiventral cartilage and parapophyses

Next, we assessed the effect of *nkx3.2* knockout on vertebrae-associated bones and cartilages in juveniles and adults. At 30 dpf mutants displayed a loss of basiventral cartilage in the precaudal rib-bearing vertebrae (Figure 7B) compared to wild-type fish (Figure 7A). At 35 dpf wild-type fish also lacked these cartilages (Figure 7C), as they have been entirely replaced through endochondral ossification by the parapophyses, small articulating bones that connect the ribs to the vertebrae. However, in both 30 dpf and 35 dpf *nkx3.2* mutants, it was clear that this endochondral ossification of basiventral cartilage had not taken place, as the parapophyses were absent (Figure 7B, D). Instead, the ribs were fused directly to the vertebrae without any articulating process, while other ribs were entirely disconnected to the vertebrae. This phenotype persisted in 90 dpf *nkx3.2* mutant adults, as seen in μCT segmented models and virtual histological sections of the vertebrae (Figure 7E-H). In contrast, the neural arches, zygapophyses, and haemal arches appeared to develop normally.

**Figure 7 –.**
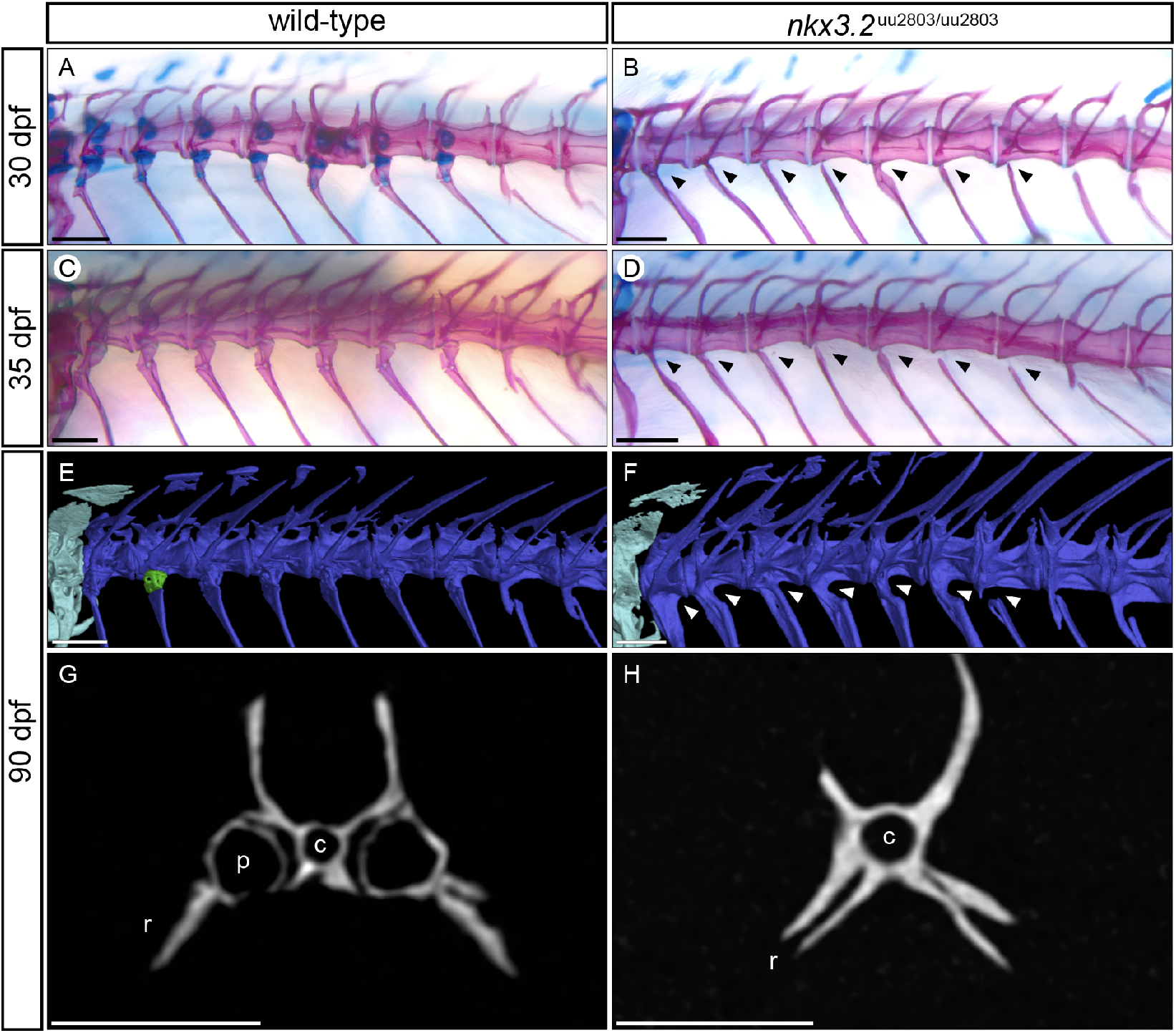
Parapophyses are absent in the rib-bearing vertebrae of *nkx3.2^uu2803/uu2803^* zebrafish. (**A-F**) Dorsolateral views of rib-bearing vertebrae in 30, 35, and 90 dpf wild-type and *nkx3.2* mutant zebrafish. (**A-D**) Cartilage- and bone-stained juvenile zebrafish, (**E, F**) μCT models. (**G, H**) μCT virtual transverse cross-sections of wild-type and *nkx3.2* mutant rib-bearing vertebrae at 90 dpf. Arrowheads in (**B, D, F**) indicate the absence of parapophyses on the rib-bearing vertebrae. A single parapophysis has been highlighted in green in (**E**) as a visual aid. c – centrum, p – parapophysis, r – rib. Scale bars: 200μm(**A-D**), 500μm (**E-H**).

### OPT, skeletal staining and μCT reveal phenotypes in the occiput and Weberian apparatus of nkx3.2^uu2803/uu2803^ zebrafish

Comparison of OPT reconstructions of 5 dpf larvae (Figure 4) suggested an effect of *nkx3.2* knockout on some of the cartilage of the otic capsule. At 30 dpf cartilage and bone staining revealed changes in the occipital bones and cervical vertebrae in *nkx3.2* mutant compared to wild-type zebrafish. More specifically, *nkx3.2* mutants displayed smaller lateral occipital fenestrae, due to changes in shapes of exoccipital and supraoccipital as well as reduced cartilages associated with forming Weberian apparatus (Figure 5G’’, H’’).

To investigate these phenotypes in greater detail, these bones were segmented from μCT scans of 90 dpf wild-type and *nkx3.2* mutant fish. In the occipital region at the back of the skull, mutants displayed a dramatic fusion between the basioccipital and exoccipital (wildtype is shown in Figure 8A, C, E, G), making it impossible to clearly demarcate these two elements, hence they are the same colour in Figure 8B, D, F, H. A central fissure in the exoccipital (Figure 8E) was also partially or completely fused in mutants (Figure 8F). As a result of the fusion of the central fissure of the exoccipital and between the exoccipital and basioccipital, the posterodorsal opening of the cavum sinus impar (csi) was lost or greatly deformed (Figure 8H). The anteroventral surface of the basioccipital that contacts the parasphenoid and prootic was highly convex in mutants, compared to only slightly convex in wild-types (Figure 8C, D; Figure S4). The posterodorsal exoccipital struts were impacted by the cervical vertebrae in *nkx3.2* mutants (Figure 8B), causing the lateral occipital fenestrae to be reduced in area and rotated posteromedially.

**Figure 8 –.**
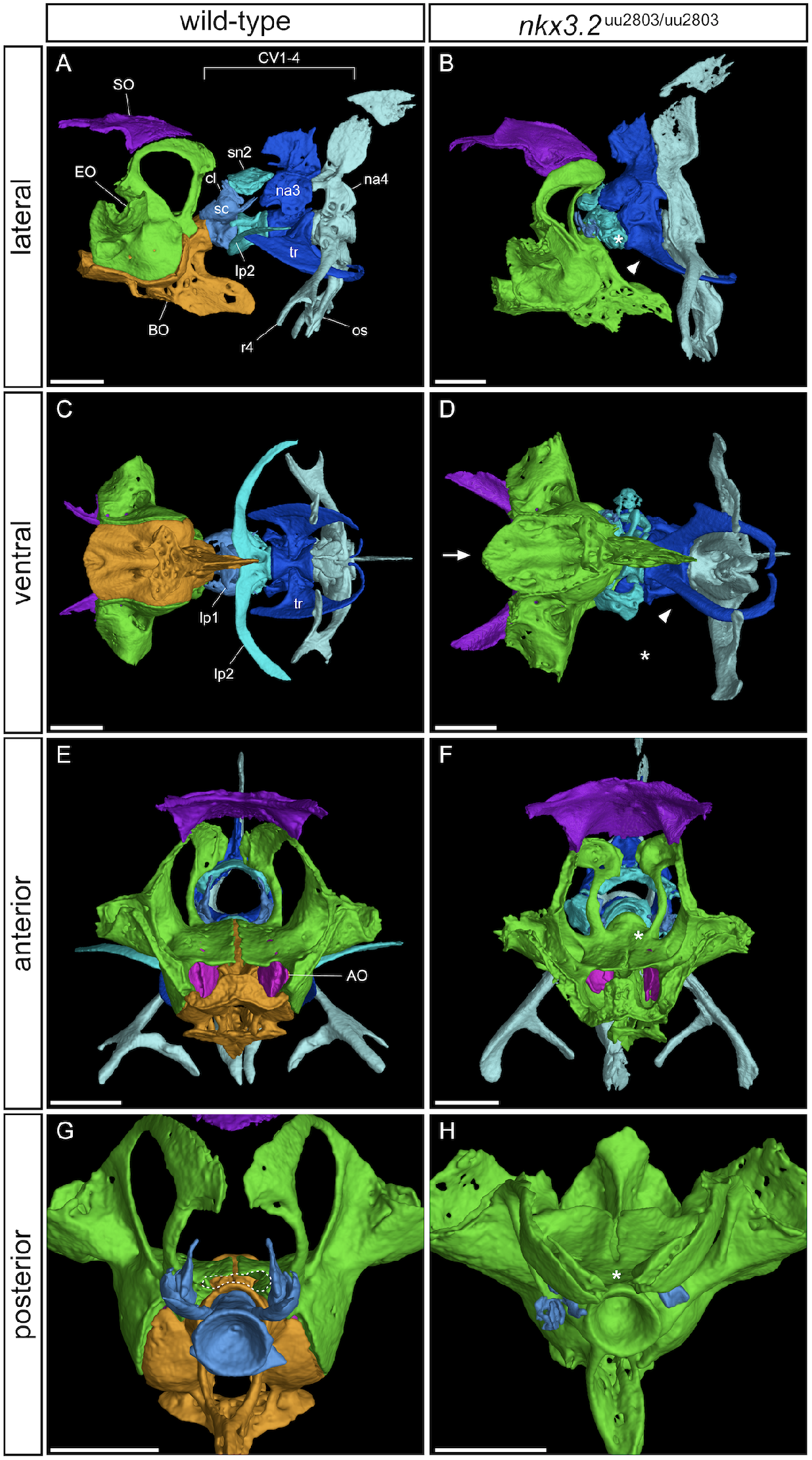
μCT reveals phenotypes in the occiput and Weberian apparatus of *nkx3.2^uu2803/uu2803^* zebrafish. Lateral, ventral, anterior, and posterior views of wild-type (**A, C, E, G**) and homozygous mutant (**B, D, F, H**) occiput and Weberian apparatus. Cervical vertebrae (CV) 2-4 have been removed in (**G, H**) for clarity. Asterisks in (**B**, **D**) indicates the absence or severe reduction of lateral process 2 from CV2, arrowheads indicate absence of anterior ramus of tripus on CV3, arrow indicates the V-shaped anteroventral edge of basipccipital. Asterisk in (**F**) indicates the posterior fusion of the dorsal fissure of the exoccipital. Dotted line in (**G**) highlights the cavum sinus impar (csi), while the asterisk in (**H**) indicates its absence. AO – asteriscus otolith, BO – basioccipital, EO – exoccipital, SO – supraoccipital, cl – claustrum, lp – lateral process, na – neural arch, os – os suspensorium, r4 – rib 4, sc – scaphium, sn2 – supraneural 2, tr – tripus. Scale bars: 500μm.

Wild-type zebrafish possess four distinct cervical vertebrae (CV), while most mutants lacked the first and most anterior CV (Figure 8A, B), with one mutant also missing CV2. Dorsal to CV1 in wild-type zebrafish lies the scaphium and claustrum, which were entirely lost or highly reduced in all mutants lacking CV1 (Figure 8B, H). The lateral process (lp2) on CV2 was significantly reduced in mutants, varying between individuals from a ~50% reduction in size to almost complete loss (Figure 8D). The tripus of CV3 was deformed in mutants – the anterior ramus and articulating process were absent (Figure 8B, D). The articulation of rib 4 and the os suspensorium to CV4 was also altered in mutants, consistent with the absence of parapophyses observed in the rib-bearing vertebrae of vertebrae 5-11 (Figure 7).

### Proximal radials in the dorsal and anal median fins are longer in nkx3.2^uu2803/uu2803^ compared to wild-type adult zebrafish

In order to assess the existence of a mutant phenotype in the dorsal and anal median fins, the lengths of the five anterior-most proximal radials (pr) were measured in μCT images of 90 dpf fish, then scaled according to the standard length of each individual to account for body size variation between individuals (Figure 9). These anterior proximal radials were chosen because the radial number is variable even between individuals of the same genotype, so a one-to-one comparison of all radials is not possible. The proximal radials in the anal fin of *nkx3.2* mutants were significantly longer than those in wild-type fish (Figure 9A). Proximal radials of the dorsal fin also tended to be longer, although didn’t always reach statistical significance (Figure 9B). These increased radial lengths may partially contribute to the dorsoventrally taller profile of *nkx3.2* mutants. In addition to the increased lengths of these radials, 40% (2/5) of 90 dpf *nkx3.2* mutant fish exhibited a deformity of pr1, such that it appeared to have two shafts (Figure 9D). No such phenotypes were observed in the paired fins or caudal fin, which appeared to develop normally, although these were more challenging to measure.

**Figure 9 –.**
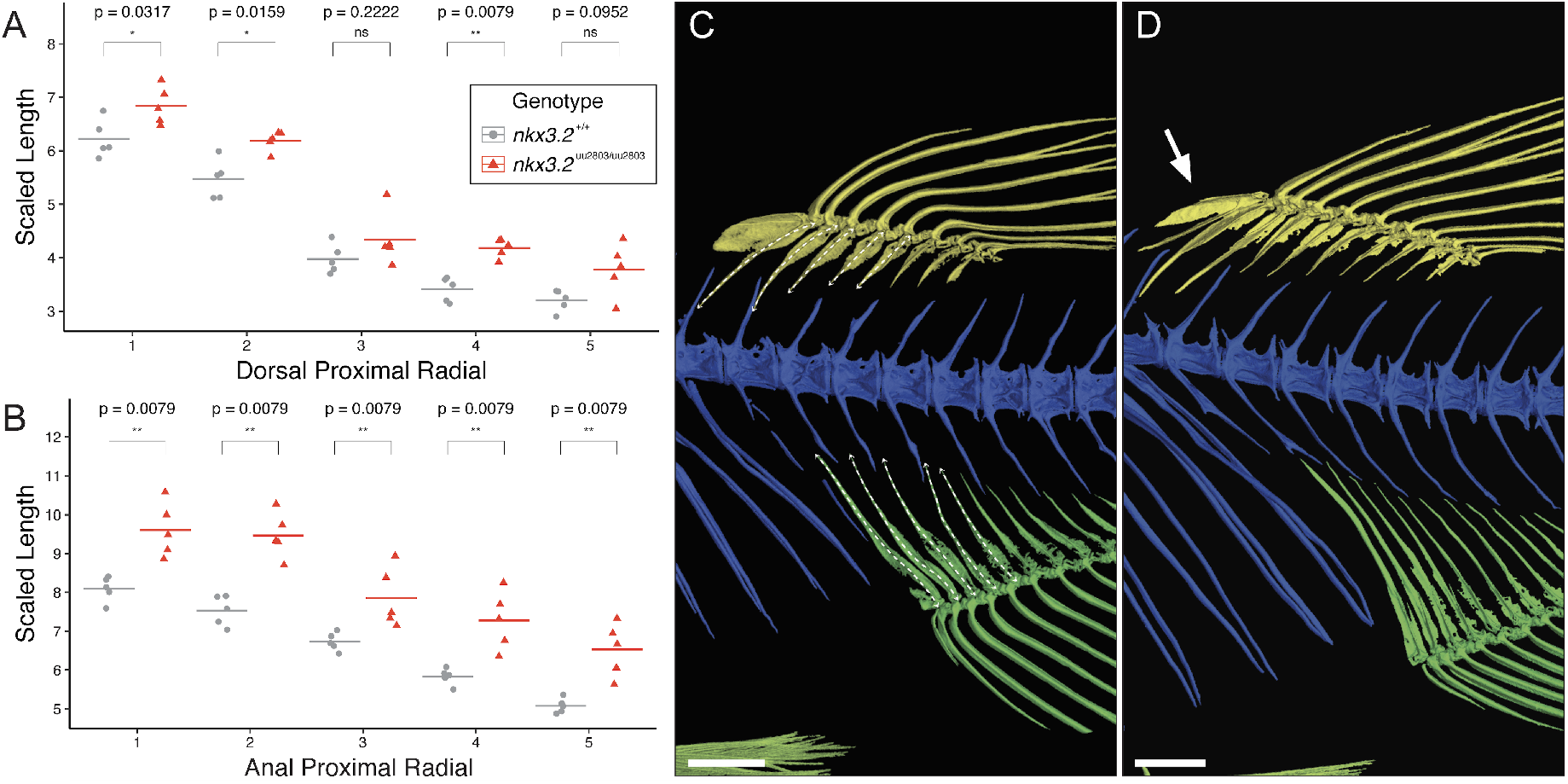
Proximal radials are longer in adult *nkx3.2^uu2803/uu2803^* zebrafish. (**A**) Proximal radials 1-5 in the dorsal fin of 90dpf *nkx.3.2* mutants are longer compared to wild-types. (**B**) Proximal radials 1-5 in the anal fin of 90dpf *nkx.3.2* mutants are substantially longer compared to wild-types. Lengths are scaled relative to standard length (SL) of each individual. Horizontal lines represent averages. (**C**) μCT segmented image 90dpf wild-type dorsal (yellow) and anal (green) fins, with white dashed arrows representing the path of the proximal radials measured. (**D**) μCT segmented image 90dpf *nkx3.2* mutant. The arrow indicates a different morphology of proximal radial 1, with the appearance of a duplicated radial shaft. Scale bars: 1mm.

## Discussion

In this study, we generated a novel zebrafish knockout mutant of *nkx3.2*. Remarkably, zebrafish homozygous for this mutation survived until adulthood, allowing us to study the mutant phenotype at a range of developmental stages up to and including adults. We employed traditional and novel techniques to characterise the effects on skeletal development this mutation caused. Consistent with previous studies, we identified a key role for nkx3.2 in the development of the jaw joint. However, we also found axial phenotypes that are novel in zebrafish, although they closely mirror phenotypes reported in human and mouse.

Homozygote *nkx3.2* mutant fish displayed a prominent open mouth phenotype from 14 dpf onwards. We did not perform experiments to study the feeding behaviour, but a recent study analysing the open mouth phenotype found that *nkx3.2* mutant fish employ ram feeding (Miyashita et al., 2020). Open mouth in nkx3.2 mutant zebrafish is caused by loss of the jaw joint and as a result, the fusion between Meckel’s and palatoquadrate cartilages of the first pharyngeal arch and loss of the retroarticular process (RAP). Our findings are consistent with the previous studies showing expression of *nkx3.2* in the jaw joint of zebrafish (Miller et al., 2003; Schwend and Ahlgren, 2009) and knockdown experiments in both zebrafish and *Xenopus* that result in the fusion of the jaw joint articulating elements Meckel’s cartilage and palatoquadrate accompanied by the loss of the retroarticular process (Lukas and Olsson, 2018a; Miller et al., 2003). These results clearly show the importance of Nkx3.2 during primary jaw joint development. Nkx3.2 loss in mouse does not affect the jaw joint but rather the associated bones of the homologous structures present in the middle ear (Tucker et al., 2004).

Our analysis of cellular organization in *nkx3.2* mutant zebrafish larvae showed that, in contrast to wild-type zebrafish, there were no jaw joint progenitor cells that could form the interzone of the jaw joint in *nkx3.2* mutant larvae. It is therefore interesting if nkx3.2 is necessary for interzone formation. Our results are consistent with findings from previous studies describing Nkx3.2 as a chondrocyte maturation inhibiting factor during skeletogenesis, which is able to repress the osteoblast maturation and skeletogenesis factor Runx2 (Lengner et al., 2005; Provot et al., 2006). As Meckel’s and palatoquadrate cartilages in *nkx3.2* mutants displayed the alignment of chondrocytes and the ability to become hypertrophic throughout the fused first arch elements. Moreover, in adult mutant zebrafish, fused articular-quadrate displayed typical characteristics of endochondrally ossified spongy bones characterized by adipocytes present in the interspace between the trabeculae (Weigele and Franz-Odendaal, 2016). It is also possible that loss of nkx3.2 could affect another transcription factor barx1. In contrast to nkx3.2, barx1 functions in repressing joint formation and promoting cartilage development. It is expressed in the first pharyngeal arch sub-intermediate domain in wild-type zebrafish, in the cartilage distal to the jaw joint (Nichols et al., 2013). Loss of nkx3.2 may allow the expansion of this expression domain, reinforcing the loss of joint identity in the region. The future examination of altered gene expression patterns of chondrogenic and joint specific factors in *nkx3.2* mutant zebrafish will be beneficial for understanding the jaw joint establishment process during development.

In addition to the primary jaw joint, *nkx3.2* is also expressed in the first arch midline domain – the symphysis joint in zebrafish embryos (Miller et al., 2003; Schwend and Ahlgren, 2009). Optical projection tomography of larvae and skeletal staining of juvenile mutant fish did not detect any abnormalities within this area except a subtle change in the shape of Meckel’s cartilage. However, our μCT analysis of the bones of adult mutant fish showed a prominent fusion of the symphysis leading to a distinctive jaw inflexibility. This disruption in symphyseal joint development is reminiscent of altered Hedgehog-signaling pathway in *con/disp1* mutant zebrafish. In these mutants *nkx3.2* expression is absent in the symphysis and reduced in the jaw joint forming region accompanied by the loss of the RAP and fusion of the symphysis (Schwend and Ahlgren, 2009). However, since we could not detect any symphysis defects in larval and juvenile *nkx3.2* mutant fish we suggest that Nkx3.2 is not essential for early symphysis development and the observed adult symphysis fusion could be a consequence of the open mouth phenotype caused by the jaw joint fusion.

In addition to confirming the essential role of Nkx3.2 in the development of the jaw joint, we describe previously unexplored axial phenotypes associated with *nkx3.2* knockout in juveniles and adults. *nkx3.2* expression in the occiput, vertebrae, and median fins has previously been described (Crotwell and Mabee, 2007), but our analysis of *nkx3.2* knockout phenotypes sheds more light on the specific axial role of this gene.

The parapophyses that articulate ribs 5-11 with the vertebral centra were absent in juvenile and adult mutants, with many ribs instead ossifying directly onto the centra, and some remaining entirely disconnected from the centra, separated by a gap where a parapophysis would have been located. The paired parapophyses normally ossify endochondrally from basiventral cartilages ventrolateral to each centrum, whereas in *nkx3.2* mutants it appears that this cartilage fails to form. This basiventral phenotype is reminiscent of the vertebral defects identified in mouse mutants, namely that the ventromedial vertebral ossification centres fail to form (Akazawa et al., 2000; Lettice et al., 1999; Tribioli and Lufkin, 1999). Similar defects in vertebral ossification have been identified in human patients suffering from SMMD, a disease caused by inactivating mutations in *NKX3.2* (Agarwal et al., 2003; Hellemans et al., 2009; Silverman and Reiley, 1985; Simon et al., 2012; Simsek-Kiper et al., 2019).

A major difference between the centra of teleosts and mammalian tetrapods is that the former directly ossify into bone without a cartilage precursor (Bensimon-Brito et al 2012), while the latter ossify endochondrally (Fleming et al, 2015). The cartilaginous vertebral precursors in tetrapods are derived from the sclerotome, while in zebrafish the notochord sheath mineralises to form chordacentra, followed by a sclerotome-derived intermembranous bone, forming autocentra (Bensimon-Brito et al., 2012; Fleming et al., 2004; Nordvik et al., 2005). Chondrichthyans and other non-tetrapod osteichthyans also form cartilaginous anlage of vertebral bodies, indicating that teleosts represent a derived condition (Criswell et al., 2017a; Peskin et al., 2020; Zhang, 2009). The basiventral and basidorsal cartilage elements in teleosts are derived also from the sclerotome (Criswell et al., 2017b; Gadow and Abbott, 1895). Our results are consistent with a role for Nkx3.2 in sclerotome-derived cartilage development in osteichthyans, rather than in vertebral development more generally. Thus, vertebrae-associated cartilages are affected in some way in the zebrafish, mouse, and human, but lead to different specific defects in teleosts compared to mammals as a result of these fundamental differences in vertebral development. Similar skeletal phenotypes are observed in knockouts of other transcription factors known to be involved in sclerotome patterning, such as Pax1 (Koseki et al., 1993; Wallin et al., 1994) and Gli2 (Mo et al., 1997).

Defects in the cervical vertebrae is another shared phenotype caused by *nkx3.2* mutations in the zebrafish, mouse, and human. In human SMMD patients, the reduced ossification of cervical vertebrae can lead to kinking of the neck (kyknodysostosis) and secondary neurological problems associated with an injured cervical cord (Simon et al., 2012). *Nkx3.2* knockout mouse embryos display a lack of chondrogenesis in the cervical vertebral bodies (Herbrand et al., 2002; Tribioli and Lufkin, 1999), and this is also likely the case in human SMMD embryos, part of the aetiology of the severe cervical defects seen postnatally. In zebrafish and other members of the teleost superorder Ostariophysi, especially otophysians, the cervical vertebrae and their associated elements have a unique structure collectively termed the Weberian apparatus (Grande and Young, 2004). This complex of bones is adapted for transmitting sound from the swim bladder to the inner ear along a chain of bony elements connected by ligaments. These bony elements represent highly derived cervical ribs and neural arches of vertebrae 1-4. In *nkx3.2* knockout zebrafish, we observed defects in all the ventral elements of the Weberian apparatus: lateral process 2, the tripus, and rib 4/os suspensorium. In the tripus and rib 4, the parts of these elements that are likely derived from basiventral cartilages – the anterior ramus of the tripus and both articulating processes – are absent or malformed such that the tripus and rib 4 are fused to cervical vertebrae 3 and 4 respectively, reminiscent of the phenotype in the other rib-bearing vertebrae as a result of the absence of parapophyses. Lateral process 2 on CV2 is absent or highly reduced in mutants. Dorsal elements of the Weberian apparatus, on the other hand, such as the supraneurals and neural arches, appear relatively unaffected. However, it is difficult to interpret the direct effects of *nkx3.2* knockout on the scaphium and claustrum because of the common loss of CV1 and the space dorsal to it that we suspect results from the impaction of the cervical vertebrae into the occiput. These results further support an essential role of Nkx3.2 that is restricted to the basiventral and not the basidorsal cartilage in zebrafish.

Basiventral cartilage elements were once thought to be a gnathostome-specific feature until it was revealed that hagfish, one of the two extant agnathan vertebrate taxa, possessed basiventral elements (Ota et al., 2011). The lamprey, on the other hand, only possesses basidorsal elements, leading to the hypothesis that this taxon secondarily lost basiventral elements and that the ancestral vertebrate possessed both basiventral and basidorsal elements (Ota et al., 2011). Studies on the involvement of Nkx3.2 in the development of hagfish basiventral elements would shed light on its potentially pivotal role in this important vertebrate innovation – was it essential in early vertebrates, or was it only recruited later in the gnathostome lineage?

In the occiput of adult *nkx3.2* mutant zebrafish, we observed a partial or complete fusion between the bones of the exoccipital and basioccipital that resulted in a partial or complete loss of the cavum sinus impar. Our OPT results revealed a subtle but significant reduction in cartilage staining intensity in this region at 5 dpf, which would have gone unnoticed comparing individual images of the larvae by eye. These results highlight the utility of the OPT method in identifying subtle phenotypes in larval stages, especially in cases where the characterisation of adult phenotypes may not be possible. In mouse knockouts, these same occipital bones are misshapen and underdeveloped (Akazawa et al., 2000; Lettice et al., 1999; Tribioli and Lufkin, 1999), although fusions between them have not been described. In addition, these *Nkx3.2* knockout mice display an absence of the supraoccipital bone, while the supraoccipital in zebrafish appears to develop normally. There are no reports of any defects to the occipital bones of human SMMD patients, although it is not clear whether this is because no defects exist or because they have been overlooked, likely a result of the difficulty in studying affected individuals *in utero* or shortly after birth.

In gnathostomes, the most anterior somites contribute to the occipital bones (Ferguson and Graham, 2004; Maddin et al., 2020; Morin-Kensicki et al., 2002). The sclerotome from these occipital somites contributes to the basioccipital and exoccipital (Couly et al., 1993; Müller and O’Rahilly, 2003, 1994), so the occipital phenotypes observed in zebrafish and mouse *nkx3.2/Nkx3.2* mutants are consistent with the role of Nkx3.2 in the sclerotome. The combination of these occipital phenotypes, particularly the defects in the cavum sinus impar and Weberian apparatus suggest that *nkx3.2* knockout zebrafish should have a severe hearing impairment, which is supported by our anecdotal observations that larval mutants fail to respond to tapping on their plates, while their heterozygous or wild-type siblings do.

*nkx3.2/Nkx3.2* expression has previously been identified in the median fins of zebrafish (Crotwell and Mabee, 2007) and the limb buds and digits of mice (Akazawa et al., 2000; Tribioli et al., 1997; Tribioli and Lufkin, 1999). We identified longer proximal radials, relative to standard body length, in adult *nkx3.2* mutant zebrafish, which is consistent with results in humans and mice. Like mammalian limb bones, teleost median proximal radials first develop as hyaline cartilage before endochondral ossification takes place (Benjamin et al., 1992; Konstantinidis and Conway, 2010). Human SMMD patients often display several limb defects postnatally: long limbs, the presence of large epiphyses and irregular metaphyses that give the disease its name, and the presence of pseudoepiphyses in the digits combined with reduced ossification of the carpals (Agarwal et al., 2003; Hellemans et al., 2009; Silverman and Reiley, 1985; Simon et al., 2012). Limb defects have not been reported in mouse mutants (Herbrand et al., 2002; Lettice et al., 1999; Tribioli and Lufkin, 1999), but changes in the regulation of *Nkx3.2* expression have been linked to tibia length (Castro et al., 2019), suggesting that the gene has a similar role in humans and mice and that the reason limb defects have not been found in mouse mutants is that the mutation is perinatally fatal, while these limb phenotypes appear postnatally (Hellemans et al., 2009). These results are consistent with a common role for Nkx3.2 in repressing chondrocyte maturation and therefore endochondral bone formation, as downregulation or gene knockout results in longer endochondral limb and median fin bones in all three species (Castro et al., 2019; Hellemans et al., 2009), and *Nkx3.2* overexpression in mice causes the opposite – skeletal dwarfism (Jeong et al., 2017).

The proximal radials of the dorsal and anal median fins develop with contributions from somite-derived cells including the sclerotome (Freitas et al., 2006; Shimada et al., 2013), while paired fins and limbs are derived from the lateral plate mesoderm (Shimada et al., 2013; Zeller et al., 2009). Even though these different skeletal structures develop from different progenitor populations, it has long been recognised that the paired limb buds redeploy developmental mechanisms that first evolved in the median fins (Freitas et al., 2006). Our results suggest that *nkx3.2* expression may have been co-opted from median fin to limb development in the sarcopterygian lineage or even during the evolution of tetrapods, as *nkx3.2* expression in actinopterygian paired fins has not been reported and we were unable to identify any phenotypes in the paired fins of adult *nkx3.2* mutant zebrafish. Definitive studies on the expression of *nkx3.2* in the fins of more basal actinopterygians such as sturgeon, gar, and bichir, in addition to chondrichthyans, are needed to provide a solid phylogenetic footing for this conclusion, as it may be that teleost paired fins represent the derived state in this regard (Davis et al., 2007).

This study highlights the role of *nkx3.2* in the development of different skeletal tissues. In the jaw and endochondral proximal radials, mutant phenotypes are consistent with a loss of chondrocyte maturation inhibition, while in the basiventral cartilages along the precaudal vertebrae, Nkx3.2 appears to be required for the onset or maintenance of cartilage development. Comparing these results with studies of amniote model systems and human disease reveal a largely consistent function of this gene between teleosts and amniotes, suggesting their inheritance from an early osteichthyan ancestor. Future studies in chondrichthyans and agnathans will further inform our understanding of the function of Nkx3.2 in early vertebrate evolution.

## Materials and Methods

### Ethical statement

All animal experimental procedures were approved by the local ethics committee for animal research in Uppsala, Sweden (permit number C161/4, 5.8.18-18096/2019). All procedures for the experiments were performed in accordance with the animal welfare guidelines of the Swedish National Board for Laboratory Animals.

### CRISPR/Cas9 target design

Two sgRNAs targeting the single zebrafish *nkx3.2* gene with no predicted off-target effects were designed using the online software CHOPCHOP (Labun et al., 2016), both targeting the first exon: 5’-GATCAGGAATCCGCGGCCAA-3’ and 5’-GTCGTTGTCCTCGCTCAGCC-3’. The second base of each target was modified to “G” in order to allow T7 transcription without modifications. The sgRNAs were prepared as previously described (Varshney et al., 2015), creating a fragment consisting of the T7 promotor, the targeted gene-specific sequence, and the guide core sequence. The sgRNAs were synthesised by in vitro transcription using the HiScribe T7 High Yield RNA Synthesis Kit (New England Biolabs, Ipswich, MA). Cas9 mRNA was prepared by in vitro transcription with the mMESSAGE mMACHINE T3 Transcription Kit (Life Technologies, Carlsbad, CA) using 500 ng of linearised plasmid that was retrieved from 5 μg of p-T3TS-nCas9n plasmid (plasmid #46757; Addgene, Cambridge, MA) digested with XbaI (New England Biolabs, Ipswich, MA). The products were purified, and their integrity was assessed using a denaturation gel.

### Generation of zebrafish mutant line

Fertilised zebrafish (*Danio rerio*) eggs were obtained by natural spawning of *Tg(sox10:egfp)* line (Carney et al., 2006). Embryos were injected at the one-cell stage with 150 pg of Cas9 mRNA and 50 pg of each sgRNA in RNase-free water as previously described (Varshney et al., 2015), and maintained at 28.5°C in E3 medium (Westerfield, 2000). The efficiency of the targets was estimated by the CRISPR-Somatic Tissue Activity Test (STAT) methodology in eight embryos at two days post-injection, as previously described (Carrington et al., 2015). The injected founder zebrafish (F0) were raised and incrossed. For genotyping the F1 zebrafish, DNA was extracted from a 1-3 mm amputation of the adult zebrafish caudal fin by lysing the tissue in 30 μl of 50 mM NaOH for 20 min at 95°C, adding 60 μl of 0.1 mM Tris and diluting the obtained material (1:10). For the initial genotyping step, FLA analysis was used. 2 μl of DNA (50-200 ng) was added to Platinum Taq DNA Polymerase. The PCR mix was incubated at 94°C for 12 min followed by 35 cycles of 94°C for 30 sec, 57°C for 30 sec, 72°C for 30 sec, and final extension at 72°C for 10 min. Size determination was carried out on a 3130XL ABI Genetic Analyzer (Applied Biosystems, Waltham, MA) and the data were analysed using the Peak Scanner Software (Thermo Fisher Scientific, Waltham, MA). For the fish that screened positive for the variant, the FLA results were confirmed by Sanger sequencing.

### Founder screening and identification of heterozygous adult fish

One strain with an allele containing a frameshift deletion resulting in a premature stop codon (*nkx3.2*^uu2803^) was selected for further experiments. The identified F1 founders were crossed with wild-type zebrafish (AB strain), and their adult offspring (F2) was genotyped. Heterozygous F2 fish of mutant line *nkx3.2*^uu2803^ were incrossed and the offspring was observed with bright-field and fluorescence microscopy. Embryos showing phenotypes were sacrificed and genotyped by FLA and/or Sanger sequencing (forward primer: 5’-TGTAAAAC-GACGGCCAGTGAGGAGTCTCGCCATCTGAA-3’; reverse primer: 5’-GTGTCTTACAGATGAA-GCTTTGAGTGGT-3’).

### In vivo microscopy

Fluorescent images were obtained with an inverted Leica TCS SP5 confocal microscope using LAS-AF software (Leica Microsystems). Embryos were sedated with 0.16% MS-222 and embedded in 0.8% low melting agarose onto the glass bottom of the 35mm dishes. To prevent drying, embedded embryos were covered with system water containing 0.16% MS-222. Screening for GFP in zebrafish larvae was performed using a Leica M205FCA fluorescence microscope with the appropriate filter.

### Histological analysis

Zebrafish juveniles and adults at 14, 30 and 90 dpf were fixed in 4% PFA and washed in PBST buffer. 30 and 90 dpf fish were decalcified in 0.5 M EDTA for one week with EDTA exchange every third day. Fish were transferred into 99.5% ethanol, followed by Xylene and embedded into paraffin. Sagittal head sections of 6μm were prepared with Leica RM2155 Rotary Microtome. Tissue sections were deparaffinised with xylene and re-hydrated through 99.5% to 70% ethanol series and transferred to water. Sections were then stained with Nuclear Fast Red (Vector Laboratories, Burlingame, CA) for 30 sec followed by a brief water rinse and dehydration in 95% and 99.9% ethanol. Prior to mounting with VectaMount (Vector Laboratories, Burlingame, CA), slides were washed in Clear-Rite 3 (Richard-Allan Scientific, Kalamazoo, MI). Sections were imaged with 40x objective on Hamamatsu NanoZoomer S60 Digital Slide Scanner.

### Skeletal staining

Staining of cartilage and bone was done based on the previously published protocol by Walker and Kimmel (2007). Zebrafish wild-type and mutant fish at 5, 9, 14 and 30 dpf were euthanised, fixed in 4% PFA and transferred to 50% ethanol. For cartilage staining, specimens were immersed in alcian blue solution (0.02% Alcian Blue 8 GX, 50mM MgCL2, 70% ethanol), and for bone staining, specimens were immersed in alizarin red solution (0.5% Alizarin Red S). For double staining of cartilage and bone, specimens were immersed in double staining solution (99% alcian blue solution, 1% alizarin red solution). After staining overnight, specimens were washed twice with 50% ethanol and then immersed in water for 2 hours before being bleached in a solution of 1.5% H202 and 1% KOH until pigmentation was removed. 30 dpf specimens were then immersed in trypsin solution (1% trypsin, 35% sodium tetraborate) for 30 minutes followed by incubation in a solution of 10% glycerol and 0.5% KOH for 1 hour. All specimens were imaged with a Leica M205FCA microscope in a solution of 50% glycerol and 0.25% KOH, followed by storage in 50% glycerol and 0.1% KOH.

### Optical Projection Tomography

A custom-built Optical Projection Tomography (OPT) system was used for imaging of the zebrafish embryos fixed at 5 dpf and stained with alcian blue (Sharpe et al., 2002; Zhang et al., 2020). The OPT system, reconstruction algorithms, and alignment workflow were based on the previously described method (Allalou et al., 2017). All embryos were kept in 99% glycerol before they were loaded into the system for imaging. The rotational images were acquired using a 3X telecentric objective with a pixel resolution of 1.15 μm/pixel. The tomographic 3D reconstruction was done using a filtered back projection (FBP) algorithm in MATLAB (Release R2015b; MathWorks, Natick, MA) together with the ASTRA Toolbox (Palenstijn *et al*., 2013). For the data alignment, the registration toolbox elastix (Klein et al., 2010; Shamonin et al., 2014) was used. To reduce the computational time all 3D volumes in the registration were down-sampled to half the resolution.

The registration workflow was similar to the methods described by Allalou *et al*. (2017) where the wildtype fish were initially aligned and used to create an average reference fish using an Iterative Shape Averaging (ISA) algorithm (Rohlfing et al., 2001). All wild-type (*n*=10) and *nkx3.2*^uu2803/uu2803^ (*n*=11) zebrafish were then aligned to the reference. After the alignment, a voxel-wise method was used to detect voxels that are significantly different between the groups. The Mann-Whitney U test was used to compare corresponding voxels in wild-type and mutant. The p-value threshold is set using a false discovery rate (FDR) (Noble, 2009) and a permutation test (Simpson et al., 2013). The FDR was set so that those random groupings showed only a small number of significant voxel differences (p<2.5×10^-4^; FDR=0.045). All registration and analysis were done on the green channel of the RGB images.

### Micro-computed tomography and segmentation

Five wild-type and five *nkx3.2*^uu2803/uu2803^ zebrafish were fixed at 90 dpf and analysed with micro-computed tomography (μCT, SkyScan 1172, Bruker microCT, Belgium) at a voltage of 60 kV, a current of 167 μA, and an isotropic voxel size of 5.43 μm. Cross-sections were reconstructed using software package NRecon (NRecon 1.6.10, Bruker microCT, Belgium). The specimens were placed in 2mL Eppendorf tubes filled with 1% agarose and the 10ul pipette tip was used to keep the mouths of some wild-type fish in the open position. BMP image stacks obtained with μCT were imported into, segmented, and imaged using VGStudio MAX version 3.2.5 (Volume Graphics, Germany).

### Statistical analyses

Proximal radial length measurements of 90dpf wild-type and *nkx3.2* mutant fish were normalised by the standard length of the fish to give a scaled length in arbitrary units. Graphs were prepared using the ggplot2 package (Wickham, 2016) in R (R Development Core Team, 2020). A Wilcoxon test was used in the analysis.

## Author Contributions

TH, LW, and JL designed this project; all authors performed experimental work and analysed the data; TH, LW, and JL wrote the paper with contributions from other authors.

## Acknowledgements

TH was supported by Vetenskapsrådet (Starting grant 621-2012-4673). The development of the OPT system was funded by a development project at SciLifeLab Uppsala (2017) and a Technology Development grant at SciLifeLab (2018) both awarded to AA. VGStudio MAX licence and some lab expenses were covered by a Wallenberg Scholarship awarded to Prof. Per E. Ahlberg, who also offered thoughtful comments on the manuscript. We thank the Genome Engineering Zebrafish facility in SciLifeLab Uppsala for generating CRISPR/Cas9 mutants, and Prof. Åsa Mackenzie for access to the Hamamatsu NanoZoomer S60 Digital Slide Scanner.

