## Supplementary Material for "The Broad Role of Nkx3.2 in the Development of the Zebrafish Axial Skeleton"

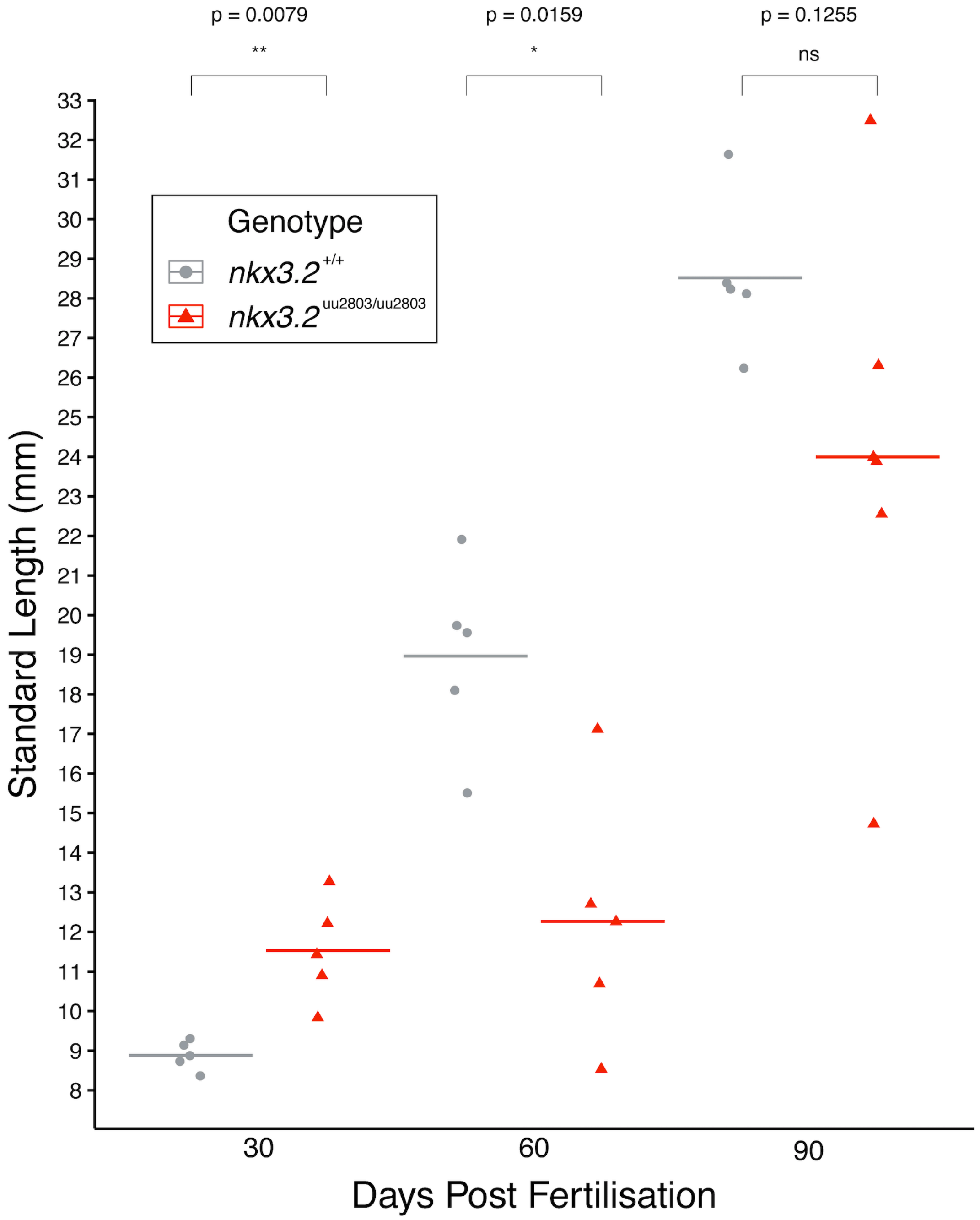


**Figure S1 – Standard lengths of a sample of wild-type and mutant zebrafish at 30, 60, and 90 dpf.** *nkx3.2* mutant zebrafish display variable body sizes at different developmental stages. The significant body size difference at 30 dpf may the result of different tank densities.

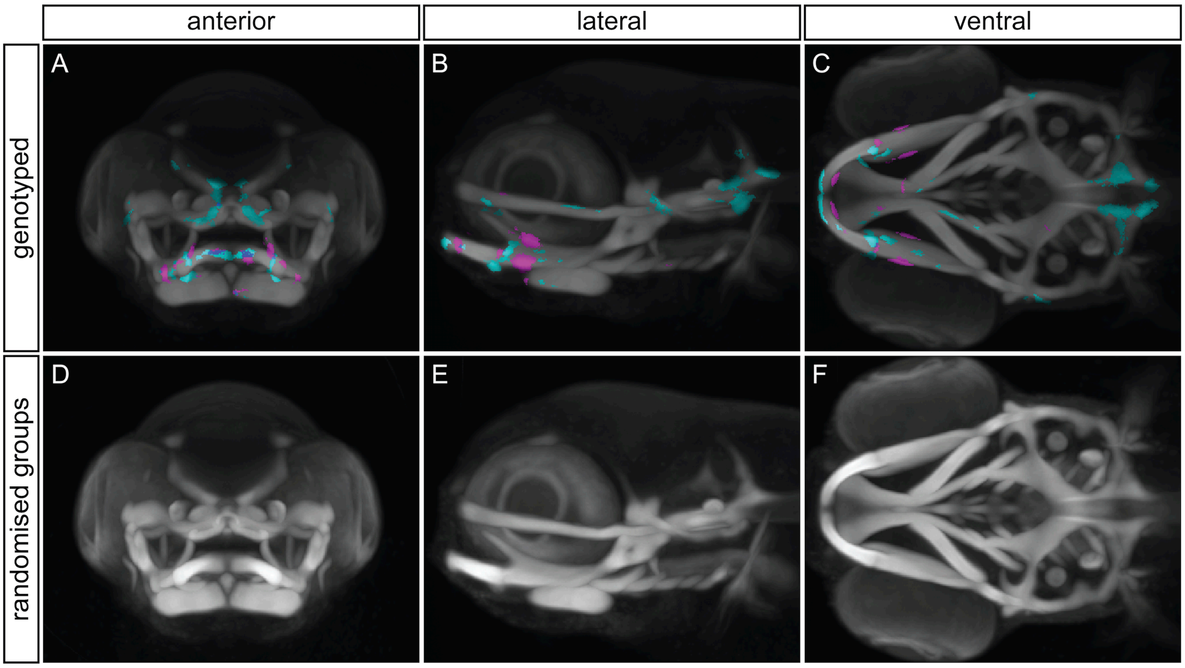


**Figure S2 – Voxel-wise method is robust to false positives.** (**A, B, C**) Maximum projection of both the 5 dpf wild-type (n=10) and *nkx3.2* knockout (n=11) groups, with coloured voxels representing voxels with statistically significant ($p<2.5x10^{-5}$) differences in intensity. Cyan shows voxels with higher intensity in wild-type group and magenta shows voxels with higher intensity in *nkx3.2* knockout group. (**D, E, F**) The same analysis performed using a randomised subset of larvae instead of comparing wild-type and *nkx3.2* knockout groups. The absence of cyan and magenta voxels indicates a lack of statistically significant false positives in the comparison of these randomised groups.

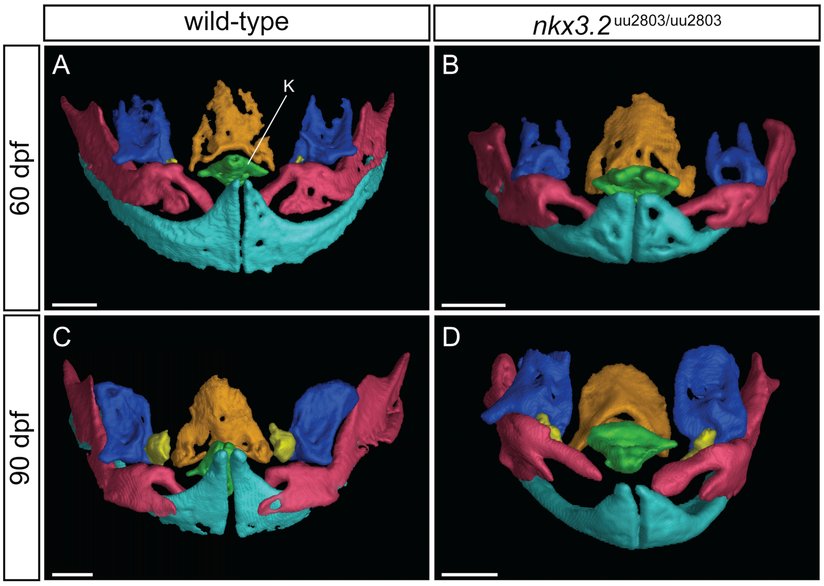


**Figure S3 – Kinethmoid bone is present in adult *nkx3.2* mutants.** (**A-D**) Dorsal view of isolated upper jaw elements – premaxilla (light blue), maxilla (pink), kinethmoid (green), preethmoid (yellow), palatine (dark blue), and ethmoid (orange). Kinethmoid is also indicated by K in (**A**). (**A**, **B**) 60 dpf wild-type (open mouth) and *nkx3.2* mutant, respectively. (**C**, **D**) 90 dpf wild-type (open mouth) and *nkx3.2* mutant, respectively. Scale bars: 200µm.

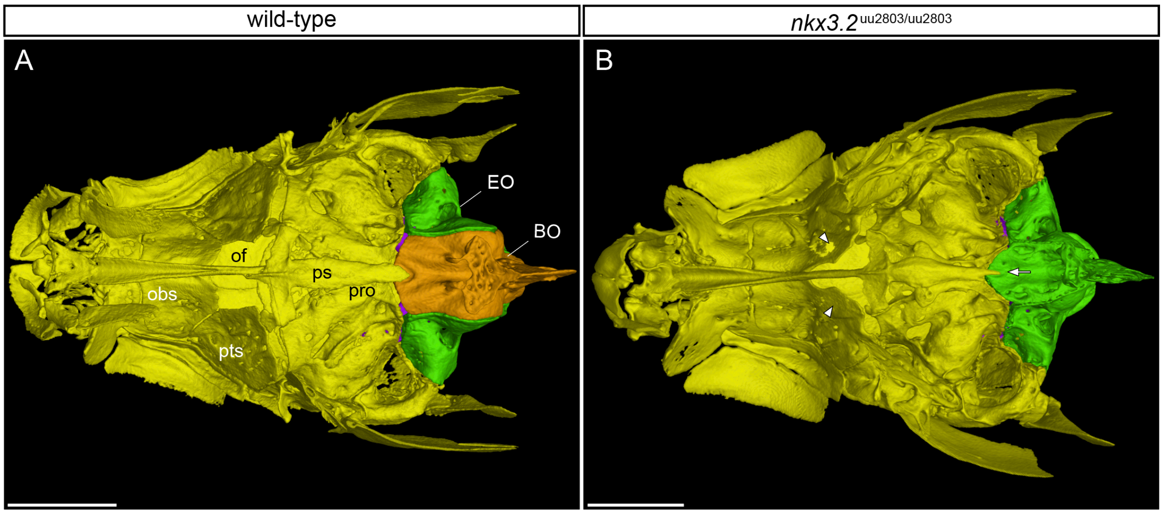


**Figure S4 – Ventral views of the adult skull.** 90dpf wild-type (**A**) and *nkx3.2* mutant (**B**). Mutants display a reduced area of the orbital fenestra as a result of posteromedial expansion of the orbitosphenoid and pterosphenoid (arrowheads). The anterior edge of the ventral surface of the basioccipital is V-shaped as it meets the parasphenoid (arrow). BO – basioccipital, EO – exoccipital, obs – orbitosphenoid, of – optic foramen, pro – prootic, ps – parasphenoid, pts – pterosphenoid. Scale bars: 1mm.

964
